# Sex-dependent effects of a glutamine supplementation on metabolic disorders, intestinal barrier function and gut microbiota in mice with diet-induced obesity

**DOI:** 10.1101/2024.11.21.624683

**Authors:** Candice Lefebvre, Adam Tiffay, Charles-Edward Breemeersch, Virginie Dreux, Christine Bôle-Feysot, Charlène Guérin, Jonathan Breton, Elise Maximin, Magali Monnoye, Pierre Déchelotte, Véronique Douard, Moïse Coëffier, Alexis Goichon

## Abstract

**Rationale:** Obesity is often associated with sex-dependent metabolic complications, in which altered intestinal barrier function and gut microbiota contribute. Glutamine (Gln) supplementation previously showed beneficial effects on gut barrier function and glycemic control. We thus aimed to characterize in mice the sex-dependent effects of a Gln supplementation during high fat diet induced obesity.

**Methods:** Male and female C57BL/6 mice received a standard (SD) or high fat diet (HFD; 60% kcal from fat) during 14 weeks (W14). From W12, mice received or not Gln in drinking water (2g/kg/day; n=12/group). Body composition, glucose tolerance, insulin sensitivity, intestinal permeability, colonic expression of 44 genes encoding factors involved in inflammatory response and gut barrier function, cecal microbiota and inflammatory/endocrine adipose response have been assessed. Data were analyzed using *t*-test or Mann-Whitney test (HFD effect), and two-way ANOVA (HFD x Gln) followed by Bonferroni post-tests.

**Results:** In both male and female mice, Gln supplementation failed to improve body weight and body composition. However, Gln reduced glucose intolerance in HFD males (AUC reduced by 14.57%, p<0.05) that was associated to a partial restoration of plasma resistin and insulin and to a trend for a limitation of adipose inflammatory response. In males, Gln did not affect gut microbiota composition and colonic response. To the opposite, in females fed HFD, Gln supplementation led to gut microbiota changes (increase of *Bacteroidota* and *Pseudomonadota* phyla; increase of *Muribaculaceae* and *Tannerellaceae* families), increased colonic inflammatory markers (TNFα, IL-1β, TLR4, Myd88, Irf3) that were associated to increased inflammatory response in subcutaneous adipose tissue and increased HOMA-IR.

**Conclusions:** High fat diet mice exhibit sex-dependent response to glutamine supplementation with protective effects in males and harmful effects in females. The role of gut microbiota should be deeply deciphered in further investigations.

## Introduction

Obesity is a major public health issue because of its prevalence and complications. In Europe countries, 50% of the adult population is overweight and obese (1). In the United States, 50% of adults will be obese by 2030 (2). Obesity is often associated with metabolic complications, e.g. type 2 diabetes, dyslipidemia, and increases the risk to develop chronic diseases, e.g. inflammatory and cardiovascular diseases, non-alcoholic fatty liver diseases, cancer. The role of gut microbiota and gut barrier function during obesity has been highlighted by regulating low-grade systemic inflammation and glycemic control related-dysregulations (3,4). A disruption of the intestinal barrier function has been described in animal models of obesity (3,5,6), even if the data in humans remain controversial (7–9). Targeting gut barrier function is a way to limit obesity-associated metabolic complications in patients with obesity. A restoration of intestinal permeability has been reported after weight loss in patients with obesity, either after severe caloric restriction (10,11) or after bariatric surgery (12). To our knowledge, the impact of GLP-1/GIP receptors agonists on gut barrier function has not yet been reported. Again, to limit the development of obesity-associated complications, other approaches (less invasive and/or low economic costs) need to be evaluate.

Glutamine (Gln), a non-essential amino acid, appears as a good candidate to limit obesity-associated metabolic complications since alterations of Gln metabolism in white adipose tissue has been reported during obesity (13). In addition, Gln regulates intestinal barrier function both during experimental and clinical researches (14). Petrus et *al.* described that changes in Gln metabolism in adipocytes contribute to modify post-translational protein modification, O-GlcNacylation, and then adipose tissue inflammation and expansion during obesity (13). Interestingly, intra-peritoneal injections of Gln were able to restore adipocyte O-GlcNacylation level. Gln supplementation was also associated with gut barrier restoration in different experimental models (15–18) but also in humans during randomized controlled trials (19,20). However, the impact of Gln supplementation on gut barrier function during obesity has not yet been reported while Gln supplementation in subjects with obesity was able to reduce fat mass and waist circumference(21–23) and to restore *Firmicutes/Bacteroideta* ratio (24). Finally, we recently reported a sex-dependent response to high fat diet (HFD) in mice (25), as also reported by others (26–28). To our knowledge, the sex-dependent response to Gln supplementation during obesity remains unknown.

Thus, in the present study, we aimed to assess the impact of oral Gln supplementation on metabolic complications in both male and female mice with diet-induced obesity as well as its effects on obesity-associated inflammatory responses and gut dysbiosis.

## Material and methods

### Animal experimentation

Six-week-old male (M) and female (F) C57BL/6J mice (Janvier Labs, Le Genest-Saint-Isle, France, n=48/sex) were acclimatized in a controlled environment (23°C with a 12 hours light-dark cycle) for 4 days with free access to water and a Standard Diet (SD, 3.34 kcal/g, 1314, Altromin, Germany). M- and F-mice were then randomized into two groups receiving either SD or High Fat Diet for 14 weeks (HFD, 5.21 kcal/g; D12492i, Research Diet Inc., US). HFD supplied 60% kcal as fat, 20% kcal as carbohydrates and 20% kcal as protein. From 12 weeks (W), mice received or not glutamine (Gln, G5792, Merck, Germany) in drinking water to provide 2 g.kg^−1^ of body weight per day^−1^. Thus, for each sex, 4 experimental groups were formed (n=12/sub-group): SD without Gln supplementation (SD), SD with Gln supplementation (SD+Gln), HFD without Gln supplementation (HFD), and HFD with Gln supplementation (HFD+Gln). Body weight was measured weekly and body composition was measured by EchoMRI (EchoMRI, Houston, US) on W4, W8, W12 and W14 of the protocol, which was approved by the regional ethics committee (authorization on APAFIS #29283-2021012114574889 v5) and performed in accordance with the current French and European regulations.

### Oral glucose (OGTT) and insulin (ITT) tolerance tests

Oral glucose (OGTT) and insulin (ITT) tolerance tests were realized before (W12) and after Gln supplementation (W14) as previously described (25). Briefly, for OGTT, after an overnight fasting followed by an oral gavage of glucose (1 g. kg^-1^ of body weight; G8769, Merck), glycaemia was measured at 0, 15, 30, 45, 60 and 90 minutes using a glucometer (Accu Check Guide, Roche, Switzerland). For ITT, after a 2 hours fasting period followed by an intraperitoneal (i.p.) injection of human insulin (0.75 U.kg^-1^ of body weight; I9278, Merck, Germany), glycaemia was measured in the same way as for the OGTT.

### Intestinal permeability

After a 2 hours fasting period and 3 hours before euthanasia, mice received a FITC-dextran solution (4 kDa, 40 mg/mL, Merck) by oral gavage at 10 µL/g of body weight to evaluate whole intestinal permeability *in vivo*, as previously described (25). Colonic paracellular permeability was also evaluated by measuring fluxes of Alexa Fluor 680-dextran (3 kDa, 0.02 mg/mL, Merck) and Lucifer Yellow (400 Da, 2.5 mg/mL, Merck) across colonic mucosa mounting in Ussing Chambers (Harvard Apparatus, US) after 3 hours at 37°C. The fluorescence intensities of FITC-dextran (excitation at 485 nm, emission at 535 nm) in plasma or Alexa Fluor 680-Dextran (excitation at 665 nm, emission at 710 nm) and Lucifer Yellow (excitation at 428 nm, emission at 540 nm) in serosal medium were measured with the Spark multimode microplate reader (Tecan, UK).

### Tissue sampling

Mice were anesthetized by i.p. injection of ketamine/xylazine solution (100 and 10 mg/kg, respectively). Blood samples were collected, centrifuged for 15 min at 3000*xg* at 4°C and plasma was frozen at -80°C. Subcutaneous and perigonadal adipose tissue depots and colon were collected, washed with ice-cold PBS, immediately frozen in liquid nitrogen and stored at -80°C until analysis. Cecal content was collected to evaluate microbiota and short-chain fatty acids.

### Plasma assays

Plasma levels of insulin, leptin, resistin and adiponectin were assessed using Bio-Plex Pro assays (BioRad Laboratories, Marnes-la-Coquette, France) and Luminex technology as previously described (25). Plasma urea, total proteins, alanine aminotransferase (ALAT), aspartate aminotransferase (ASAT), gamma-glutamyltransferase (GGT), total cholesterol and triglycerides (TG) concentrations were determined using a Catalyst One device (IDEXX Laboratories, Westbrook, ME, US).

### Colonic RNA extraction and RT-qPCR

Extraction of total colonic mRNAs, quantification and reverse transcription were performed as previously described (25). qPCRs were realized by using the SYBR Green technology on a QuantStudio 12K Flex real-time PCR system (Life Technologies, Carlsbad, CA, US) targeting 44 markers of interest mainly involved in inflammatory response and gut barrier function. The mean of ACTB, β2M and GAPDH genes were used as reference. Gene-specific primers were previously detailed (25).

### Short chain fatty acid analysis

Twenty to fifty grams of frozen caecal contents were used to analyzed the short-chain fatty acids (SCFAs) composition using gas chromatography (Agilent 7890B gas chromatograph, Agilent Technologies, Les Ulis, France) as previously described (25).

### Cecal microbiota analysis

Total DNA was extracted from cecal content samples with PowerFecal Pro DNA Kit (Qiagen, France) and using the mechanical bead-beating disruption method. The V3–V4 hypervariable region of the bacterial 16S rDNA was amplified by PCR (PCR1F_460: CTTTCCCTACACGACGCTCTTCCGATCTACGGRAGGCAGCAG and PCR1R_460: GGAGTTCAGACGTGTGCTCTTCCGATCTTACCAGGGTATCTAATCCT) using Phanta Max Super-Fidelity DNA Polymerase (Vazyme, PRC) (95°C for 3 min and then 35 cycles at 95°C for 15 sec, 65°C for 15 sec, and 72°C for 1 min before a final step at 72°C for 5 min). The purified amplicons were sequenced using Miseq sequencing technology (Illumina) at the GeT-PLaGe platform (Toulouse, France). The sequences were first prepared using R combined with dada2 and FROGS as previously described (25). The resulting sequences were saved in a sequence table using makeSequenceTable. The guidelines of FROGS v4.0.1 were used to detected and removed chimeras using the vsearch tool. ASVs with global abundance lower than 0.005 % were removed from the following analysis with FROGS filters and were then assigned to species using FROGS affiliation with 16S REFseq Bacteria. Subsequent analysis were done using the phyloseq R packages (29). Samples were rarefied to even sampling depths before computing within-samples compositional diversities (observed richnessn, Chao1 and Shannon index) and between-samples compositional diversity (Bray-Curtis) Raw, unrarefied OTU counts were used to produce relative abundance graphs and to find genra with significantly different abundances between SD, SD+Gln, HFD and HFD+Gln groups in male and female mice.

### Adipose tissue RNA extraction and RT-qPCR

Extraction of perigonadal and subcutaneous white adipose tissue (WAT) mRNAs, quantification and reverse transcription were performed as previously described (25). qPCRs were realized by using the SYBR Green technology on a Bio-Rad CFX96 real-time PCR system (Bio-Rad Laboratories). The mean of β2M and RPS18 was used as reference. Gene-specific primers were previously detailed (25).

### Statistical Analysis

Data were analyzed with GraphPad Prism 8.3 (GraphPad Software Inc., San Diego, CA, USA) and expressed as the mean ± standard error of the mean (SEM). To compare body weight or glycaemia changes, two-way ANOVA test were performed (TimexHFD) followed by Bonferroni post-tests. HFD and Gln supplementation effects were also determined by two-way ANOVA test followed by Bonferroni post-tests. Alpha diversity index (Observed species, and Shannon) were analyzed using 1-way ANOVA test. Phylum and family relative abundances were compared using a Mann-Whitney test. DESeq2 was used to estimate abundance fold changes between SD / SD+Gln or HFD / HFD+Gln in female mice. The ASVs were selected based on adjusted p-value (<0.05) and prevalence (more than 50 copies per sample in at least 25% of the samples).

## Results

### Impact of oral Gln supplementation in male mice

Fifteen days of Gln supplementation affected neither body weight nor fat mass gain in both SD and HFD male mice (Fig. 1A). Similarly, fasting glycaemia remained unchanged (Fig. 1B). Before Gln supplementation at W12, HFD mice exhibited a glucose intolerance that was partly restored in Gln-supplemented mice (Fig. 1C). The area under the curve (AUC) of glucose was reduced by 14.6% in Gln-supplemented HFD mice (Fig. 1C, p<0.05). By contrast, the ITT revealed that Gln supplementation did not modify insulin sensitivity (Fig. 1D). Plasma insulin remained unchanged, as well as HOMA-IR (Fig. 2). Gln did not significantly change the plasma metabolic markers of dyslipidemia (total cholesterol, triglycerides) and liver function (Supplemental Fig. S1). The plasma level of leptin was not affected by Gln supplementation in both SD and HFD male mice, whereas the significant increase of plasma resistin and adiponectin observed in HFD fed mice was partly blunted by Gln supplementation (Fig. 2). We then evaluated the mRNA profile of perigonadal and subcutaneous WAT. In perigonadal WAT, Gln supplementation did not significantly modify the mRNA levels of pro-inflammatory cytokines and of adipokines. By contrast, in the subcutaneous WAT, Gln supplementation significantly reduced IL6 mRNA level (Fig. 3). We also observed a trend for a decrease of Ccl2 mRNA level (p=0.057), encoding for MCP-1 and an interaction effect between HFD and Gln for resistin mRNA level (Fig. 3).

**Fig. 1:**
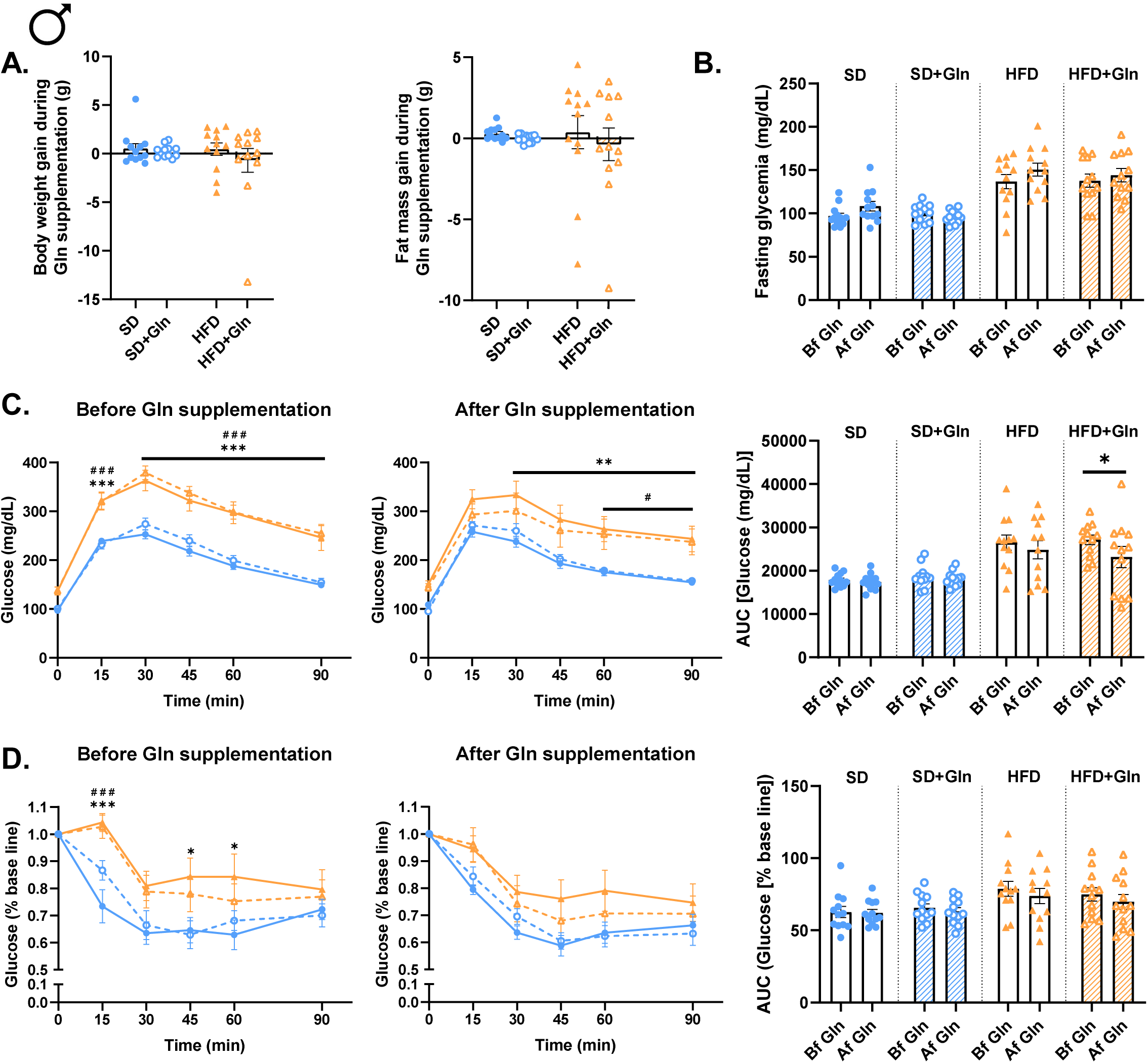
Effects of glutamine (Gln) supplementation on body weight, fat mass, glycemic control and insulin resistance in male mice fed with a standard (SD) or high fat diet (HFD). Body weight, fat mass (**A.**), fasting glycaemia (**B.**), oral glucose tolerance test (OGTT, **C.**) and insulin tolerance test (ITT, **D.**) were assessed at week 12 (W12) and week 14 (W14) in male mice fed with standard diet (SD) or high fat diet (HFD) and receiving or not glutamine (Gln) in drinking water from W12 to W14. Data are expressed as mean ± sem (n=12/group). \**p*<0.05, \*\**p*<0.01, \*\*\**p*<0.001 HFD *vs* SD, #*p*<0.05, ###*p*<0.001 HFD+Gln *vs* SD. AUC, Area under the curve.

**Fig. 2:**
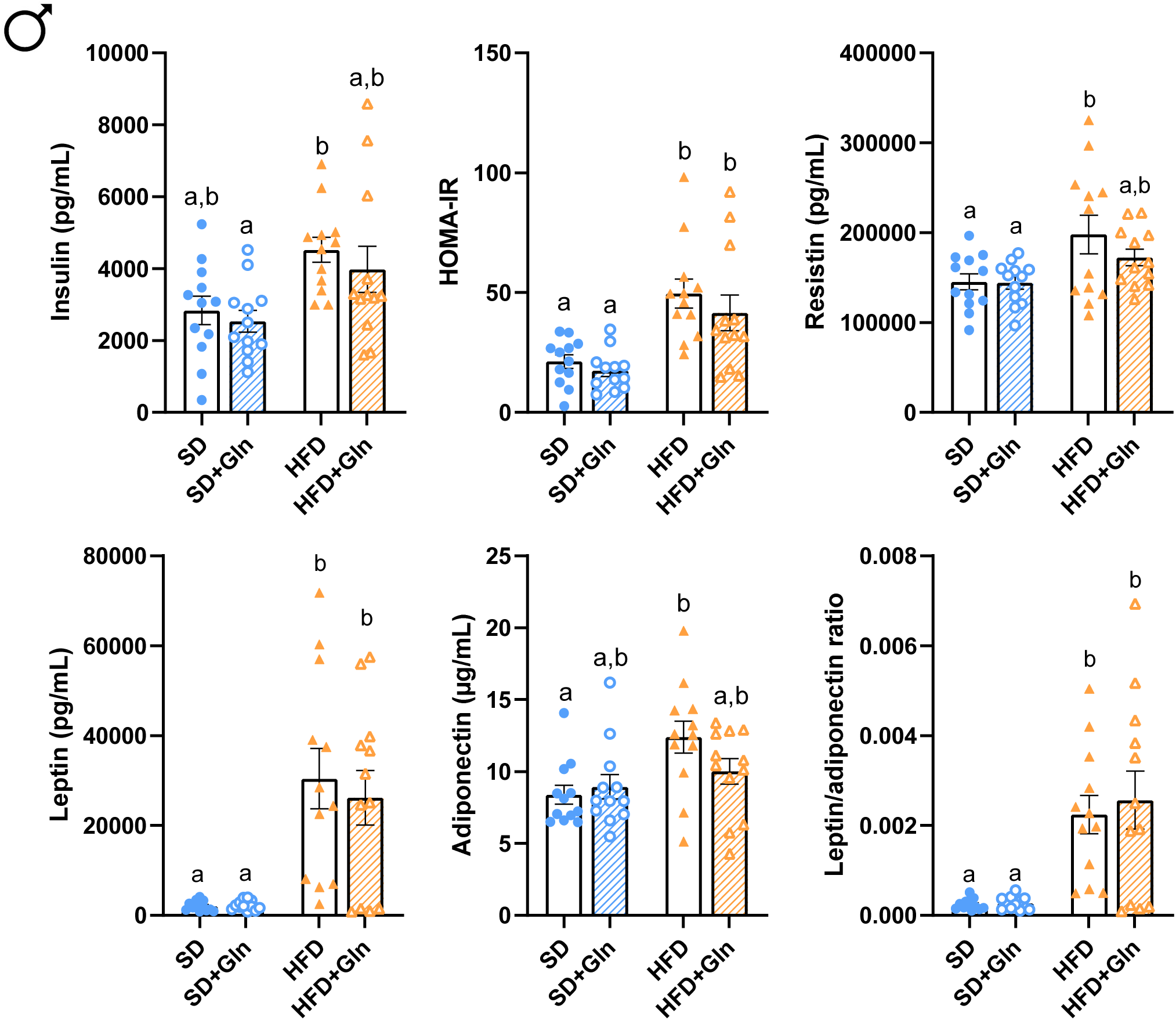
Effects of glutamine (Gln) supplementation on insulinemia, HOMA-IR and plasma adipokine levels in male mice fed with a standard (SD) or high fat diet (HFD). Plasma insulin, resistin, leptin and adiponectin concentrations were assessed at week 14 (W14) in male mice fed with standard diet (SD) or high fat diet (HFD) and receiving or not glutamine (Gln) in drinking water from W12 to W14. The HOMA-IR index was determined by the following equation: fasting serum insulin (μU/mL)LJ×LJfasting plasma glucose (mmol/L)/22.5. Data are expressed as mean ± sem (n=12/group). Values without a common letter differ significantly (p < 0.05).

**Fig. 3:**
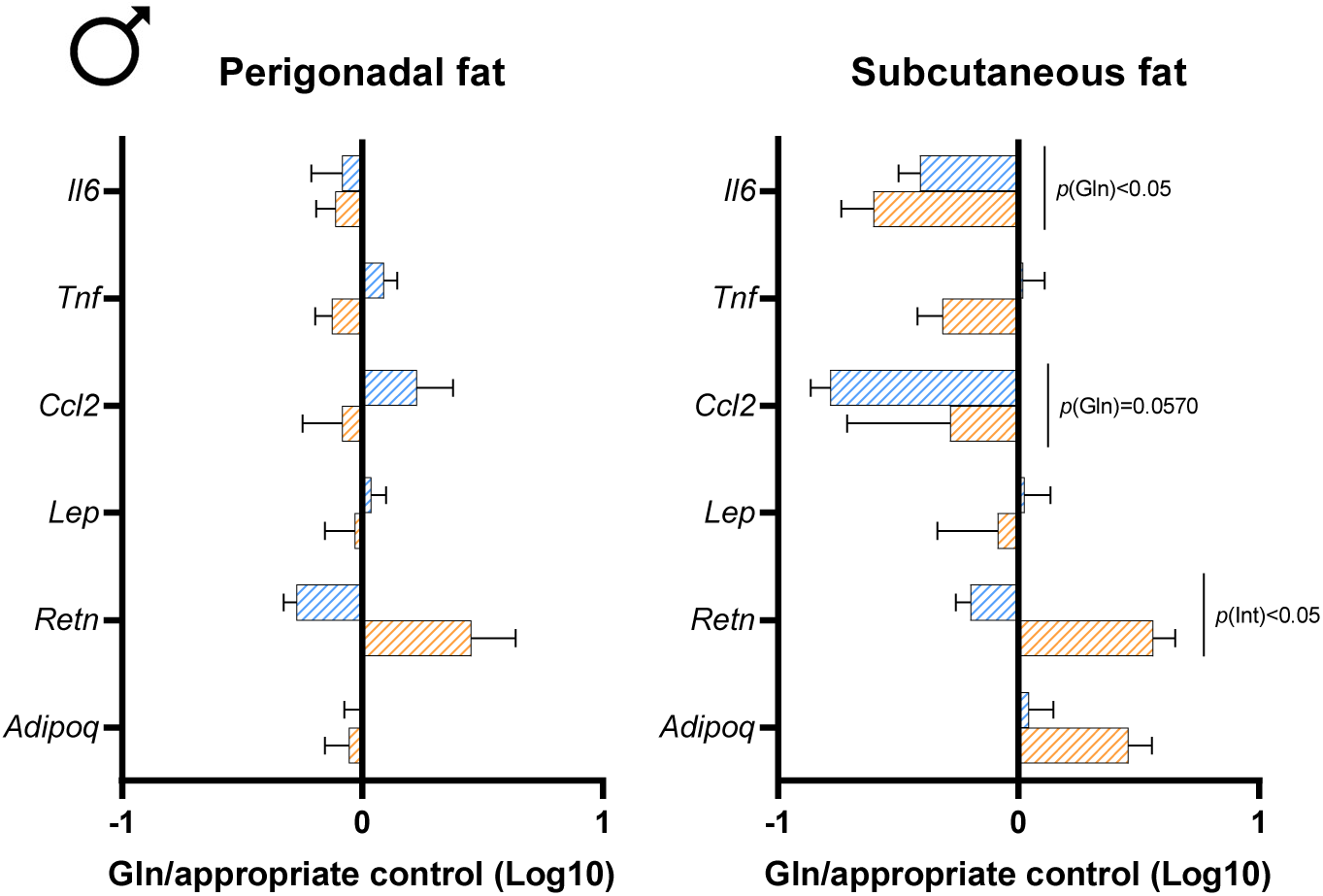
Effects of glutamine (Gln) supplementation on white adipose tissue in male mice fed with a standard (SD) or high fat diet (HFD). The levels of mRNA encoding for *Il6* (Interleukin 6), *Tnf* (Tumor necrosis factor alpha), *Ccl2* (C-C motif Chemokine ligand 2 or MCP1), *Lep* (Leptin), *Retn* (Resistin) and *Adipoq* (adiponectin) were measured in perigonadal and subcutaneous fat at week 14 (W14) in male mice fed with standard diet (SD) or high fat diet (HFD) and receiving or not glutamine (Gln) in drinking water from W12 to W14. Results were compared by two-way ANOVA (diet x Gln). Data are presented as ratio log10 [SD+Gln / SD] (blue bars) or log 10 [HFD+Gln / HFD] (Orange bars) and expressed as mean ± sem (n=12/group). * p<0.05 *vs* the appropriate controls (HFD or SD).

We then focused on the gut response since Gln is well known to regulate intestinal cell metabolism, gut barrier function and inflammatory response (14). In our study, Gln supplementation did not affect global intestinal permeability evaluated by FITC-D gavage and colonic permeability measured in Ussing chambers (Fig. 4A). A transcriptomic analysis targeting 44 genes of interest in the colonic mucosa revealed that 15 days of Gln supplementation in HFD male mice did not completely change the colonic mRNA profile expression. We only observed modifications of mRNA levels for some proteins involved in intercellular junctions: a weak reduction of Marveld2 mRNA encoding the intracellular protein named tricellulin and an increase of the mRNA level for transmembrane proteins, claudin-11 and cingulin (Fig. 4B). Interestingly, HFD male mice also exhibited a reduction of colonic mRNA level for TLR2 in response to Gln supplementation (Fig. 4C). Concerning the analysis of cecal microbiota, neither the alpha diversity (Fig. 5A) nor the phyla (Fig. 5B) and family abundance (Supplemental Fig. S2A) were significantly modified by Gln in both SD and HFD male mice. Only the relative abundance of *Prevotellaceae* was significantly decreased in response to Gln supplementation in male mice on a SD diet (Fig. 5C). Similarly, the cecal content of SCFA was not significantly changed by Gln supplementation (Fig. 5D).

**Fig. 4:**
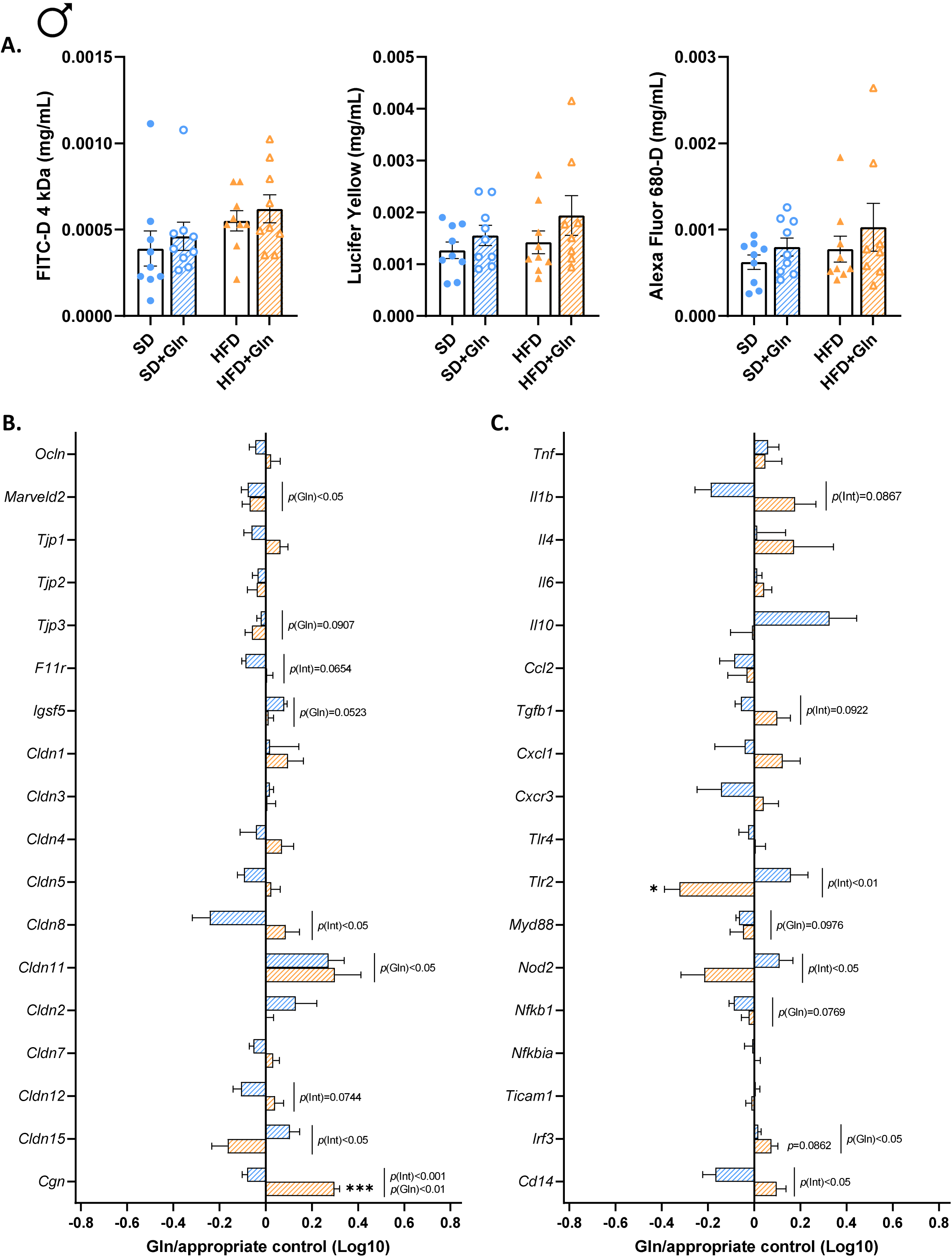
Effect of glutamine (Gln) supplementation on intestinal permeability, gut barrier and inflammatory markers in male mice fed with a standard (SD) or high fat diet (HFD). (**A.**) Global intestinal permeability (FITC-D), colonic permeability (Lucifer Yellow and Alexa Fluor 680-D) and mRNA colonic levels of gut barrier (**B.**) and inflammatory (**C.**) markers were assessed at week 14 (W14) in male mice fed with standard diet (SD) or high fat diet (HFD) and receiving or not glutamine (Gln) in drinking water from W12 to W14. Results were compared by two-way ANOVA (diet x Gln). Data are presented as ratio log10 [SD+Gln / SD] (blue bars) or log10 [HFD+Gln / HFD] (Orange bars) and expressed as mean ± sem (n=12/group). * p<0.05, *** p<0.001*vs* the appropriate controls (HFD or SD). *Ccl2*: C-C motif Chemokine ligand 2 (or MCP-1), *Cd14*: CD14 antigen, *Cgn*: Cingulin, *Cldn1*: Claudin 1, *Cldn2*: Claudin 2, *Cldn3*: Claudin 3, *Cldn4*: Claudin 4, *Cldn5*: Claudin 5, *Cldn7*: Claudin 7, *Cldn8*: Claudin 8, *Cldn11*: Claudin 11, *Cldn12*: Claudin 12, *Cldn15*: Claudin 15, *Cxcl1*: C-X-C motif chemokine ligand 1, *Cxcr3*: C-X-C motif chemokine receptor 3, *F11r*: F11 receptor or JAM-A, *Igsf5*: Immunoglobulin superfamily member 5 or JAM-4, *Il1b* :Interleukin 1 beta, *Il10*: Interleukin 10, *Il4*: Interleukin 4, *Il6* :Interleukin 6, *Irf3*: Interferon regulatory factor 3, *Marveld2*: Marvel domain-containing protein 2 or Tricellulin, *Myd88*: Myeloid differentiation primary response 88, *Nfkb1*: Nuclear factor kappa B1, *Nfkbia*: Nuclear factor kappa B inhibitor alpha, *Nod2*: Nucleotide-binding oligomerization domain-containing protein 2, *Ocln*: Occludin, *Tgfb1*: Transforming growth factor beta 1, *Ticam1*: TIR domain-containing adapter molecule 1, *Tjp1*: Tight junction protein 1 or ZO-1, *Tjp2*: Tight junction protein 2 or ZO-2; *Tjp3*: Tight junction protein 3 or ZO-3, *TLR2*: Toll-like receptor 2, *TLR4*: Toll-like receptor 4, *Tnf*: Tumor necrosis factor alpha.

**Fig. 5:**
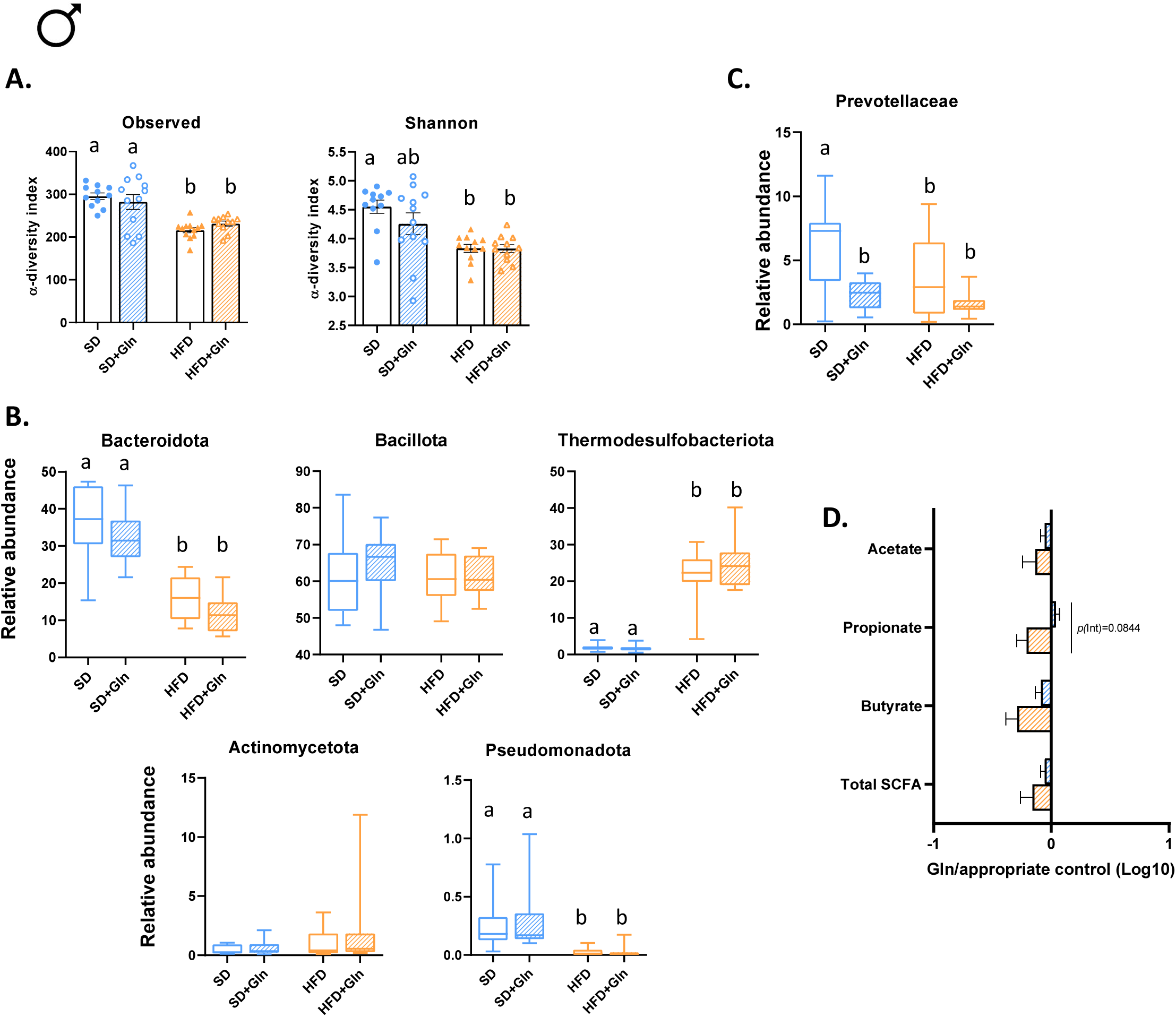
Effects of glutamine (Gln) supplementation on cecal microbiota composition and short-chain fatty acids levels (AGCC) in male mice fed with a standard (SD) or high fat diet (HFD). (**A.**) The cecal bacterial α-diversity (observed species and Shannon index), (**B.**) microbiota composition at the level of major phyla, (**C.**) family and (**D.**) AGCC levels were assessed at week 14 (W14) in male mice fed with standard diet (SD) or high fat diet (HFD) and receiving or not glutamine (Gln) in drinking water from W12 to W14. For **A.**, data are expressed as mean ± sem (n=12/group). For **B.** and **C.**, data are expressed as median and min-to-max values (n = 10-12). For **D.** data are presented as ratio log10 [SD+Gln / SD] (blue bars) or log10 [HFD+Gln / HFD] (Orange bars) and expressed as mean ± sem (n=12/group). Values without a common letter differ significantly (p<0.05).

Thus, 15 days of Gln supplementation did not have major effects on cecal microbiota, intestinal response in both SD and HFD male mice but reduced glucose intolerance and inflammatory markers in the subcutaneous adipose tissue. All these parameters have also been studied in SD and HFD female mice.

### Impact of oral Gln supplementation in female mice

In female mice, Gln supplementation did not affect body weight and fat mass (Fig. 6A), as observed in male mice. However, Gln supplementation was associated with an increase of fasting glycaemia in HFD mice (Fig. 6B) that was not observed in males, and did not reduce glucose intolerance (Fig. 6C). Insulin plasma level was not changed after Gln supplementation (Supplemental Fig. S3). As fasting glycaemia increased without modification of insulin, the HOMA-IR showed a significant increase after Gln supplementation in HFD mice (Supplemental Fig. S3), even if the ITT did not reveal any changes (Fig. 6D). Plasma lipidic and liver markers remained unchanged (Supplemental Fig. S4). While Gln supplementation did not modify the adipokine plasma concentrations (Supplemental Fig. S3), the mRNA levels for inflammatory markers and adipokines were affected mainly in the subcutaneous WAT. Indeed, by contrast to the data observed in males, the mRNA encoding for Ccl2, leptin and adiponectin were increased by Gln in the subcutaneous WAT of HFD female mice. No significant effects of Gln have been noticed in the perigonadal fat (Fig. 7).

**Fig. 6:**
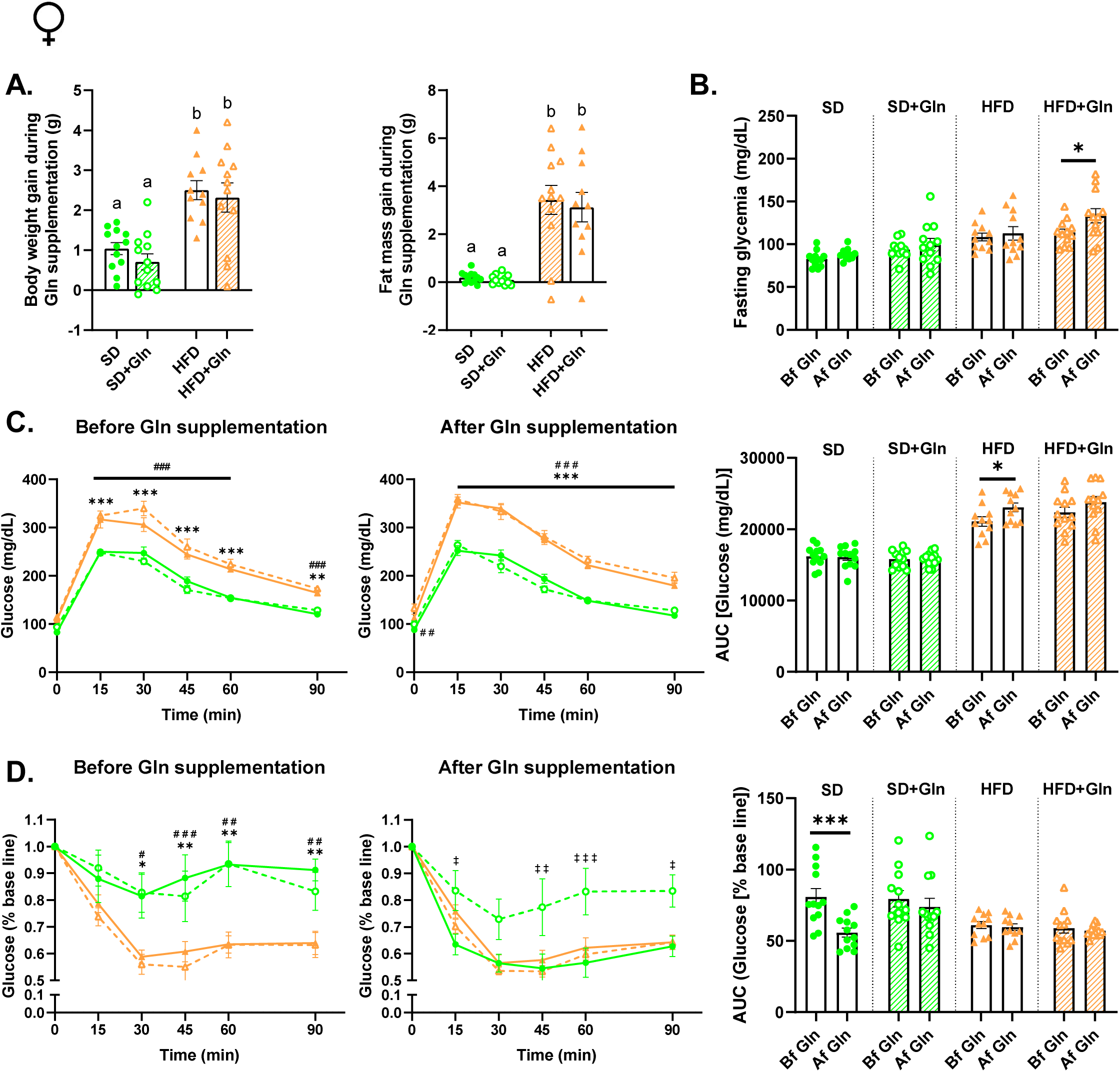
Effects of glutamine (Gln) supplementation on body weight, fat mass, glycemic control and insulin resistance in female mice fed with a standard (SD) or high fat diet (HFD). Body weight, fat mass (**A.**), fasting glycaemia (**B.**), oral glucose tolerance test (OGTT, **C.**) and insulin tolerance test (ITT, **D.**) were assessed at week 12 (W12) and week 14 (W14) in female mice fed with standard diet (SD) or high fat diet (HFD) and receiving or not glutamine (Gln) in drinking water from W12 to W14. Data are expressed as mean ± sem (n=12/group). Values without a common letter differ significantly (p<0.05). * p<0.05, ** p<0.01, *** p<0.001 HFD *vs* SD. ^‡^ p<0.05, ^‡‡^ p<0.01, ^‡‡‡^ p<0.001 SD+Gln *vs* SD. ^##^ p<0.01, ^###^ p<0.001 HFD+Gln *vs* SD. AUC, Area under the curve.

**Fig. 7:**
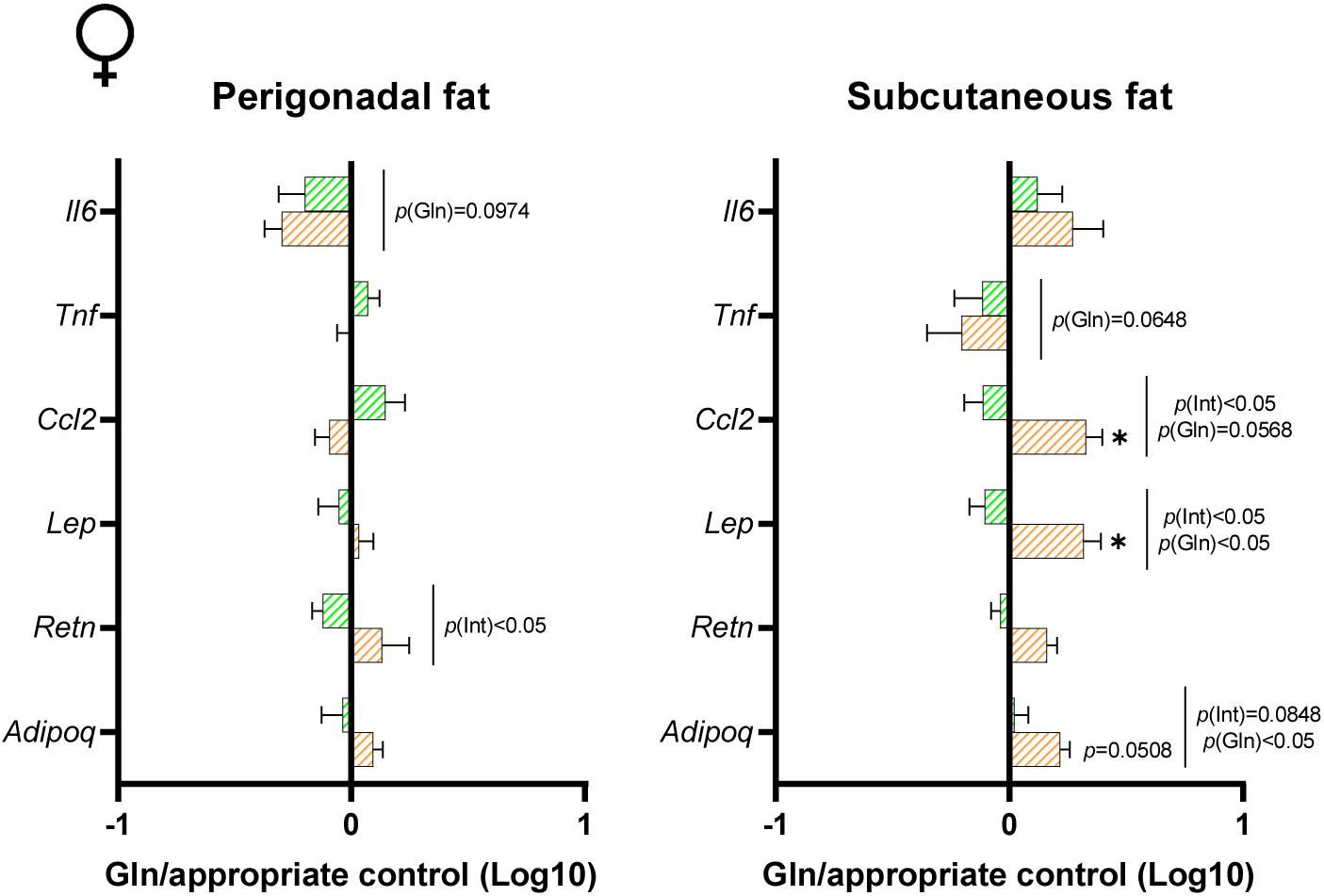
Effects of glutamine (Gln) supplementation on white adipose tissue in female mice fed with a standard (SD) or high fat diet (HFD). The levels of mRNA encoding for *Il6* (Interleukin 6), *Tnf* (Tumor necrosis factor alpha), *Ccl2* (C-C motif Chemokine ligand 2 or MCP1), *Lep* (Leptin), *Retn* (Resistin) and *Adipoq* (adiponectin) were measured in perigonadal and subcutaneous fat at week 14 (W14) in female mice fed with standard diet (SD) or high fat diet (HFD) and receiving or not glutamine (Gln) in drinking water from W12 to W14. Results were compared by two-way ANOVA (diet x Gln). Data are presented as ratio log10 [SD+Gln / SD] (green bars) or log10 [HFD+Gln / HFD] (Orange bars) and expressed as mean ± sem (n=12/group). * p<0.05 *vs* the appropriate controls (HFD or SD).

As described in our previous paper (25), HFD female mice exhibited an increase of global intestinal permeability that was however not influenced by Gln supplementation (Fig. 8A). In contrast to male mice, Gln supplementation induced several changes in the transcriptomic profile in the colonic mucosa of female mice both in SD and HFD conditions. Indeed, in SD female mice, we observed an increase of Marveld2 (tricellulin), Tjp2 (ZO-2), Igsf5 (JAM-4) mRNA expression (Fig. 8B) and a decrease of Ccl2 mRNA level (Fig. 8C). In HFD mice, Gln supplementation reduced the mRNA levels for claudin-1 and claudin-3, and increased the mRNA levels for cingulin, IL1β, TLR4, Myd88 and Irf3 (Fig. 8B, C). In addition, Gln also affected Tjp1 (ZO-1), claudin-4, claudin-15, TNFα, TLR2, Ticam1 and Cd14 mRNA levels (two-way ANOVA p-value (Gln) <0.05). Interestingly, female mice also exhibited some alterations of cecal microbiota in response to Gln supplementation that were not observed in males. Even if alpha diversity was not affected by Gln in both SD and HFD mice (Fig. 9A), we observed differences in the phyla or family abundance (Fig. 9B, C and Supplemental Fig. S2B). Indeed, Gln-supplemented HFD mice showed a higher relative abundance in *Bacteroidota* and *Pseudomonadota* compared to HFD mice (Fig. 9B). This difference was not present between SD and Gln-supplemented SD mice. Particularly, within the *Bacteroidota* phylum, we observed that the *Muribaculaceae* and *Tannerellaceae* abundances were increased in HFD mice supplemented with Gln compared to HFD mice (Fig. 9C). In Gln-supplemented SD mice, we also observed that the abundance of *Lactobacillaceae* was decreased while *Odoribacteraceae* abundance increased (Fig. 9C). In female mice, 27 and 5 ASVs were respectively modified after Gln supplementation in SD and HFD conditions. The Supplemental Fig S5 displayed the ASVs with a Log2(foldchange) > 2. Interestingly, among ASVs affected by Gln, the members of the *Lachnospiraceae, Muribaculaceae* and *Oscillospiraceae* families are mainly represented in both SD and HFD conditions. Finally, a trend for an increase of total SCFA cecal content was observed in Gln-supplemented SD mice (p=0.0531). The difference reached significance only for propionate (Fig. 9D).

**Fig. 8:**
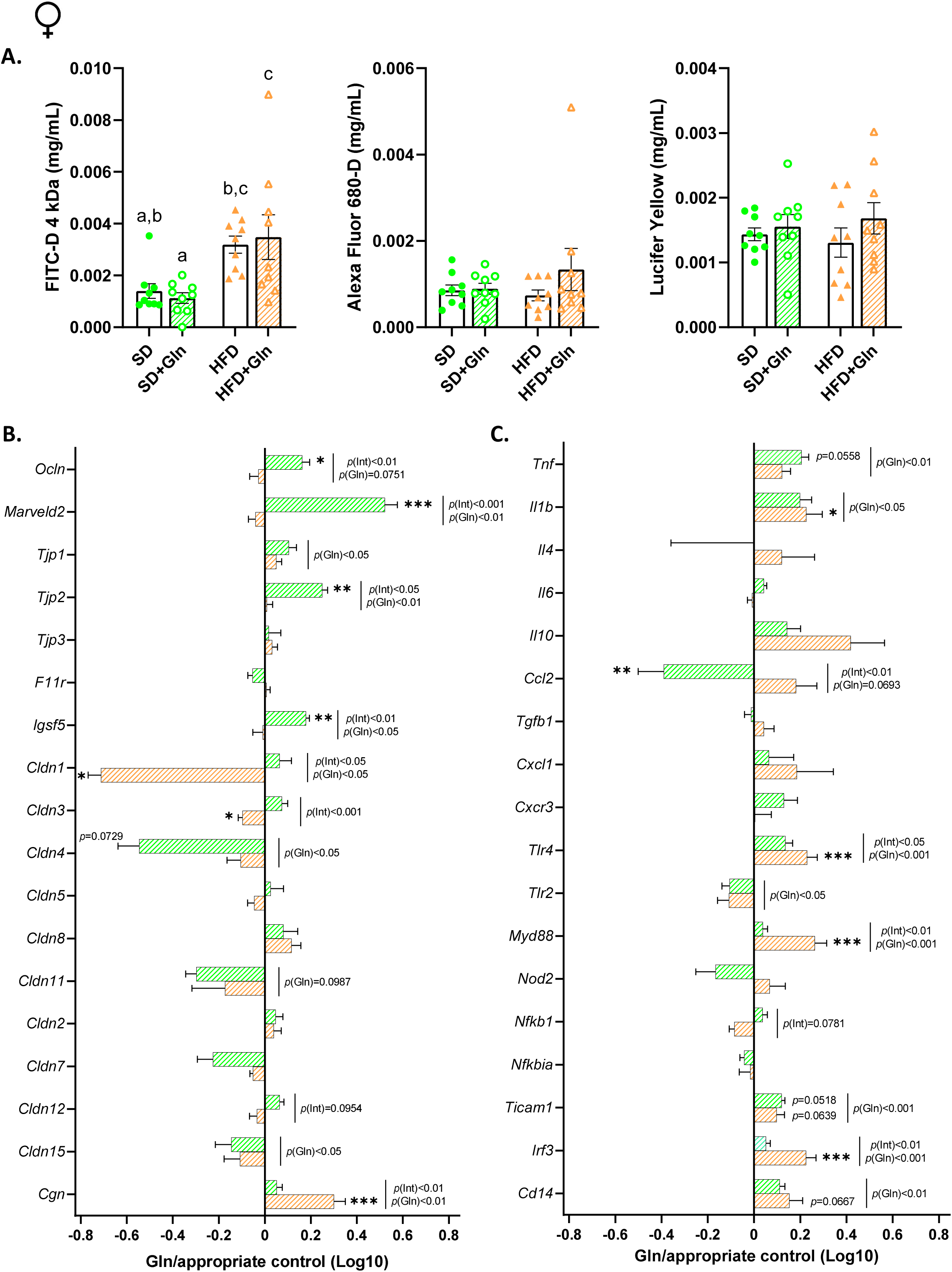
Effect of glutamine (Gln) supplementation on intestinal permeability, gut barrier and inflammatory markers in female mice fed with a standard (SD) or high fat diet (HFD). (**A.**) Global intestinal permeability (FITC-D), colonic permeability (Lucifer Yellow and Alexa Fluor 680-D) and mRNA colonic levels of gut barrier (**B.**) and inflammatory (**C.**) markers were assessed at week 14 (W14) in female mice fed with standard diet (SD) or high fat diet (HFD) and receiving or not glutamine (Gln) in drinking water from W12 to W14. Results were compared by two-way ANOVA (diet x Gln). Data are presented as ratio log10 [SD+Gln / SD] (green bars) or lo10 [HFD+Gln / HFD] (Orange bars) and expressed as mean ± sem (n=12/group). * p<0.05, ** p<0.01, *** p<0.001*vs* the appropriate controls (HFD or SD). *Ccl2*: C-C motif Chemokine ligand 2 or MCP-1, *Cd14*: CD14 antigen, *Cgn*: Cingulin, *Cldn1*: Claudin 1, *Cldn2*: Claudin 2, *Cldn3*: Claudin 3, *Cldn4*: Claudin 4, *Cldn5*: Claudin 5, *Cldn7*: Claudin 7, *Cldn8*: Claudin 8, *Cldn11*: Claudin 11, *Cldn12*: Claudin 12, *Cldn15*: Claudin 15, *Cxcl1*: C-X-C motif chemokine ligand 1, *Cxcr3*: C-X-C motif chemokine receptor 3, *F11r*: F11 receptor or JAM-A, *Igsf5*: Immunoglobulin superfamily member 5 or JAM-4, *Il1b* :Interleukin 1 beta, *Il10*: Interleukin 10, *Il4*: Interleukin 4, *Il6* :Interleukin 6, *Irf3*: Interferon regulatory factor 3, *Marveld2*: Marvel domain-containing protein 2 or Tricellulin, *Myd88*: Myeloid differentiation primary response 88, *Nfkb1*: Nuclear factor kappa B1, *Nfkbia*: Nuclear factor kappa B inhibitor alpha, *Nod2*: Nucleotide-binding oligomerization domain-containing protein 2, *Ocln*: Occludin, *Tgfb1*: Transforming growth factor beta 1, *Ticam1*: TIR domain-containing adapter molecule 1, *Tjp1*: Tight junction protein 1 or ZO-1, *Tjp2*: Tight junction protein 2 or ZO-2; *Tjp3*: Tight junction protein 3 or ZO-3, *TLR2*: Toll-like receptor 2, *TLR4*: Toll-like receptor 4, *Tnf*: Tumor necrosis factor alpha.

**Fig. 9:**
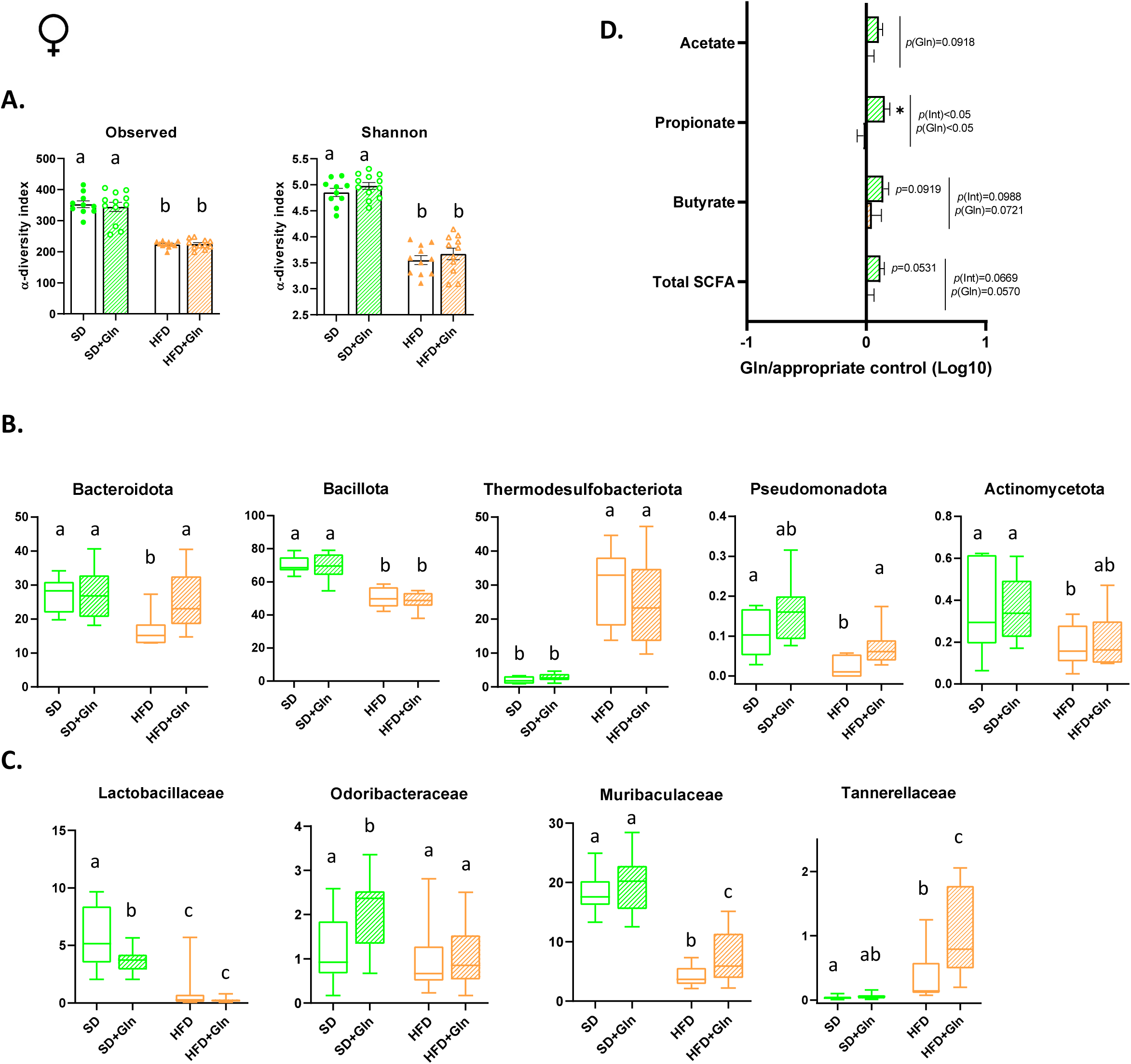
Effects of glutamine (Gln) supplementation on cecal microbiota composition and short-chain fatty acids levels (AGCC) in female mice fed with a standard (SD) or high fat diet (HFD). (**A.**) The cecal bacterial α-diversity (observed species and Shannon index), (**B.**) microbiota composition at the level of major phyla (**B.**) and families (**C.**), and AGCC levels (**D.**) were assessed at week 14 (W14) in female mice fed with standard diet (SD) or high fat diet (HFD) and receiving or not glutamine (Gln) in drinking water from W12 to W14. For **A.**, data are expressed as mean ± sem (n=10-12/group). For **B.** and **C.**, data are expressed as median and min-to-max values (n = 10-12). Results were compared by one-way ANOVA and values without a common letter significantly differ (p<0.05). For **D.**, data are presented as ratio log10 [SD+Gln / SD] (green bars) or log10 [HFD+Gln / HFD] (orange bars) and expressed as mean ± sem (n=12/group). *, p<0.05 *vs* the appropriate controls (HFD or SD).

## Discussion

Obesity and its associated co-morbidities is a major issue for public health. The role of gut microbiota and gut barrier function has been highlighted in the onset of metabolic complications by favoring a low-grade inflammatory state (3). Targeting gut microbiota and gut barrier function could be thus an interesting therapeutic strategy to limit these complications (30). Experimental data and some clinical trials showed that Gln is able to reduce intestinal hyperpermeability. So, we performed the present work to evaluate the properties of Gln to limit glucose intolerance and insulin resistance in a model of diet induced obesity in mice. Interestingly, our study reveals for the first time a sex-dependent response to Gln supplementation. Indeed, we report that oral Gln supplementation improved glycemic control in obese male mice without affecting gut microbiota composition, while in female mice, Gln supplementation altered gut microbiota composition and inflammatory response in colon and adipose tissue and seemed enhanced insulin resistance.

According with the different outcomes between men and women concerning obesity and metabolic complications, the sex-dependent response to a high fat diet in mouse has been previously reported by us (25) and others (27,34,35). Indeed, in a previous paper, we showed different kinetics in the onset of metabolic complications between male and female mice that could be related to specific alterations of gut microbiota and intestinal responses (25). In the present paper, to our knowledge, we report for the first time sex-dependent effects of Gln. Previous experimental data showed that Gln is able to restore or limit intestinal hyperpermeability *in vitro* (15,34) and *in vivo* (16,17,31). In patients with irritable bowel syndrome, oral Gln supplementation during 8 weeks was associated with a restoration of intestinal permeability measured by Lactulose/Mannitol test (19). In addition, beneficial effects of glutamine on glycemic control in intensive care patients have been observed (32,33). In the context of obesity, positive effects of Gln supplementation on body weight and waist circumference were suggested in pilot studies in humans (22,23). Gln was also able to restore the *Firmicutes/Bacteroidota* in patients with obesity (24), while glutamate/Gln fecal content tended to be reduced in patients with obesity (34). In obese rats, Abboud et al described a reduction of insulin resistance after Gln supplementation (23). Interestingly, Petrus et al highlighted a reduced Gln content in adipocytes that contributes to increase adipose inflammatory response. This was blunted by intra-peritoneal injections of Gln (13). However, it is of interest to note that clinical studies mainly enrolled women (13,22,24), whereas experimental data in rodents used exclusively males (13,23).

In the present study, Gln have protective effects in male mice by decreasing glucose intolerance that was associated with a partial restoration of fasting insulin and of adipokines, i.e. resistin and adiponectin. This was associated with a trend for a decrease of inflammatory response in the subcutaneous white adipose tissue. However, Gln failed to affect body weight, gut microbiota composition and colonic inflammatory response. To the opposite, Gln supplementation was associated with harmful effects in female mice: increase of fasting glycaemia and HOMA-IR, increase of inflammatory response in the subcutaneous adipose tissue. The role of adipose inflammatory response in the onset of insulin resistance is well established (38). The differential impact of Gln on glycemic control between males and females may be related to the sex-dependent effects of Gln on adipose tissue: anti-inflammatory effects in males, pro-inflammatory effects in females. Petrus et al reported beneficial effects of intra-peritoneal injections of Gln on adipocyte response through the regulation of O-GlcNAcylation however, only male mice were studied (13). Thus, it should be interesting to evaluate in further experiments the impact of oral Gln supplementation on O-GlcNAcylation in the subcutaneous adipose tissue according to the sex.

Interestingly, Gln supplementation was associated with modifications in the gut microbiota composition and increased colonic inflammatory response only in females. These alterations may play a role in the negative impact of Gln on adipocytes and glycemic control in females but this should be further investigated. Gut microbiota modifications by Gln were already shown in other pathophysiological conditions (35–37). In women with obesity, De Souza et al previously reported that Gln restored *Firmicutes/Bacteroidota* ratio with an increase of *Bacteroidota* from 43.2 to 52.4% of gut microbiota after Gln supplementation (24). In our study, Gln also increased *Bacteroidota* in HFD female mice but not in SD mice. At the family level, we particularly observed, in HFD fed mice, an increase of *Muribaculaceae and Tannerellaceae*, for which beneficial properties have been previously described. For instance, beneficial effects of *Lactiplantibacillus plantarum* (38) or *Lactobacillus paracasei* (39) have been associated to increase of *Muribaculaceae* that consume mucus-derived monosaccharides (40). Interestingly, a sex-dependent impact of inulin on *Muribaculaceae* during obesity has been reported. In a first study performed in male mice, inulin accelerated weight loss but did not affect *Muribaculaceae* abundance (41). In a second study, the same authors reported different effects of inulin in female mice with an increase of *Muribaculaceae* abundance (42). Finally, we observed that Gln supplementation in HFD female mice was associated with an increase of 4 ASVs among which *Lachnospiraceae, Muribaculaceae* and *Oscillospiraceae* are represented. Interestingly, a recent study showed the negative correlation between *Lachnospiraceae* and *Oscillospiraceae* abundances and body mass index (BMI) (43). Increased abundance of *Lachnospiraceae* and *Oscillospiraceae* was also associated to the beneficial effects of fucoxanthin extracts in HFD fed male mice (44) as well as the effects of inulin associated to an increase of *Muribaculaceae* in HFD fed female mice (42). However, *Lachnospiraceae* are important producers of SCFA that remained unaffected by Gln supplementation in the present study. The role of gut microbiota in the sex-dependent response to Gln needs to be further explored.

In the present study, Gln was supplied in the drinking water for two weeks in mice after 12 weeks of HFD. In Petrus study, Gln was administered by intra peritoneal injections for 14 days after 5-4 weeks of HFD (13). In Abboud study, Gln was supplied for 4 weeks by combining Gln in the drinking water and Gln administration by oral gavages 3 days per week (23). Thus, in our study, Gln supplementation was realized later than in previous works (13,23) and thus probably in more complicated animals. In addition, the oral route appears more potentially applicable for clinical use. The sex-dependent effects of Gln to prevent metabolic complications should be further evaluated.

In conclusions, we demonstrate for the first time sex-dependent effects of glutamine supplementation with protective effects on glycemic control in males but harmful effects in females that could be related to gut microbiota alterations and associated inflammatory responses. Further experiments should evaluate whether gut microbiota is responsible of the sex-dependent response to glutamine in obese mice.

### Ethics statement

The protocol was approved by the regional ethics committee (authorization on APAFIS #29283-2021012114574889 v5) and performed in accordance with the current French and European regulations.

## Acknowledgements

We thank Pamela Lecras and Dr David Vaudry from the PRIMACEN platform (HeRaCleS Inserm US51, CNRS UAR 2026, Université de Rouen Normandie, France) and the staff of GenoToul platform (Toulouse, France).

## Author’s contribution

Conceptualization, CL, AG, PD, VDo and MC; CL, AT, EM, MM, AG and MC, formal analysis; CL, AT, CE-B, VDr, CBF, CG, JB, MM, VDo and AG, investigation; CL, VDo, AG and MC, original draft preparation; writing review and editing, AG, VDo and MC.

## Figures legends

**Supplemental Fig. S1:**
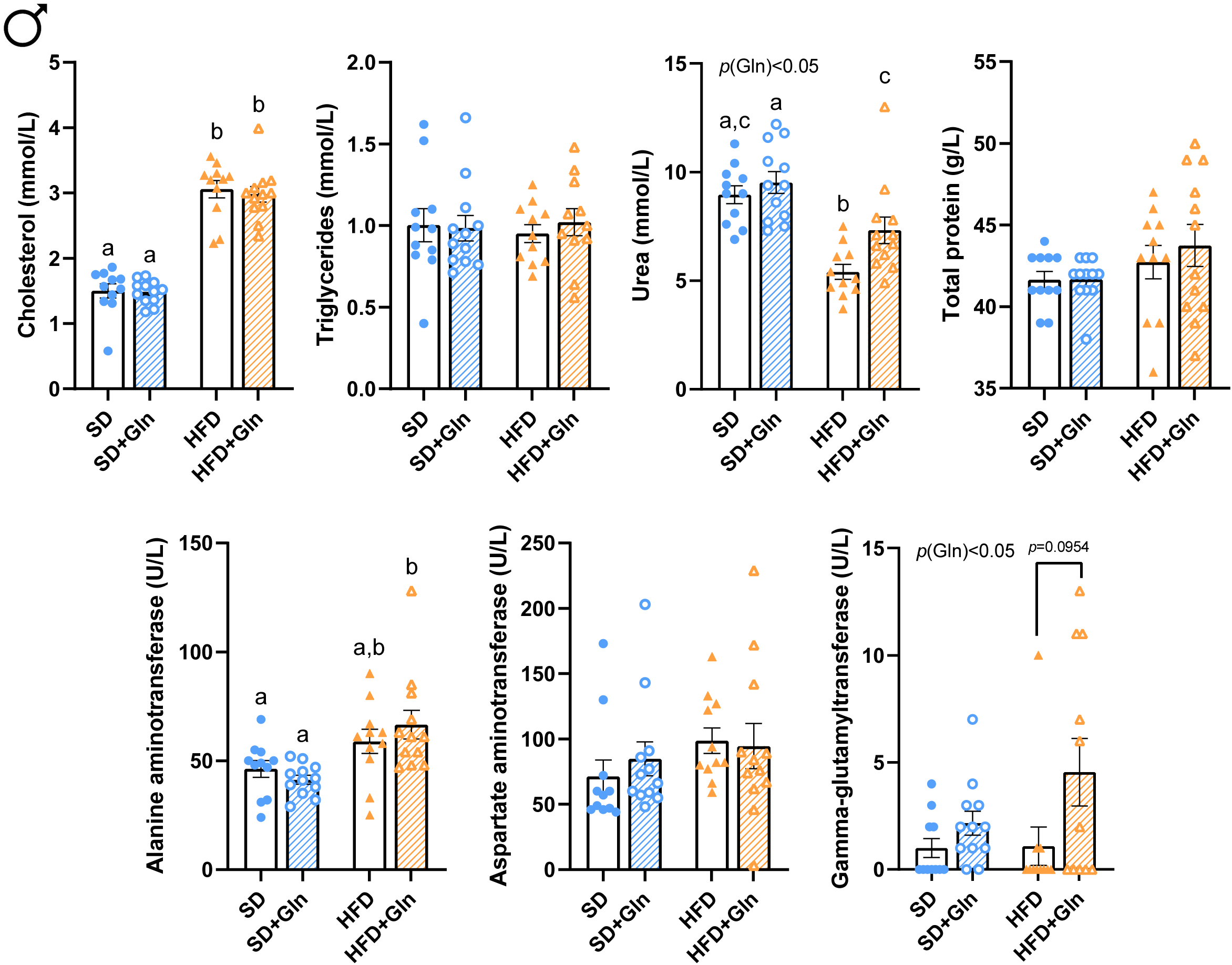
Effects of glutamine (Gln) supplementation on metabolic and liver plasma markers in male mice fed with a standard (SD) or high fat diet (HFD). Cholesterol, triglycerides, urea, total proteins, alanine aminotransferase, aspartate aminotransferase and gamma-glutamyltransferase plasma concentrations were assessed at week 14 (W14) in male mice fed with standard diet (SD) or high fat diet (HFD) and receiving or not glutamine (Gln) in drinking water from W12 to W14. Data are expressed as mean ± sem (n=12/group). Values without a common letter differ significantly (p < 0.05).

**Supplemental Fig. S2:**
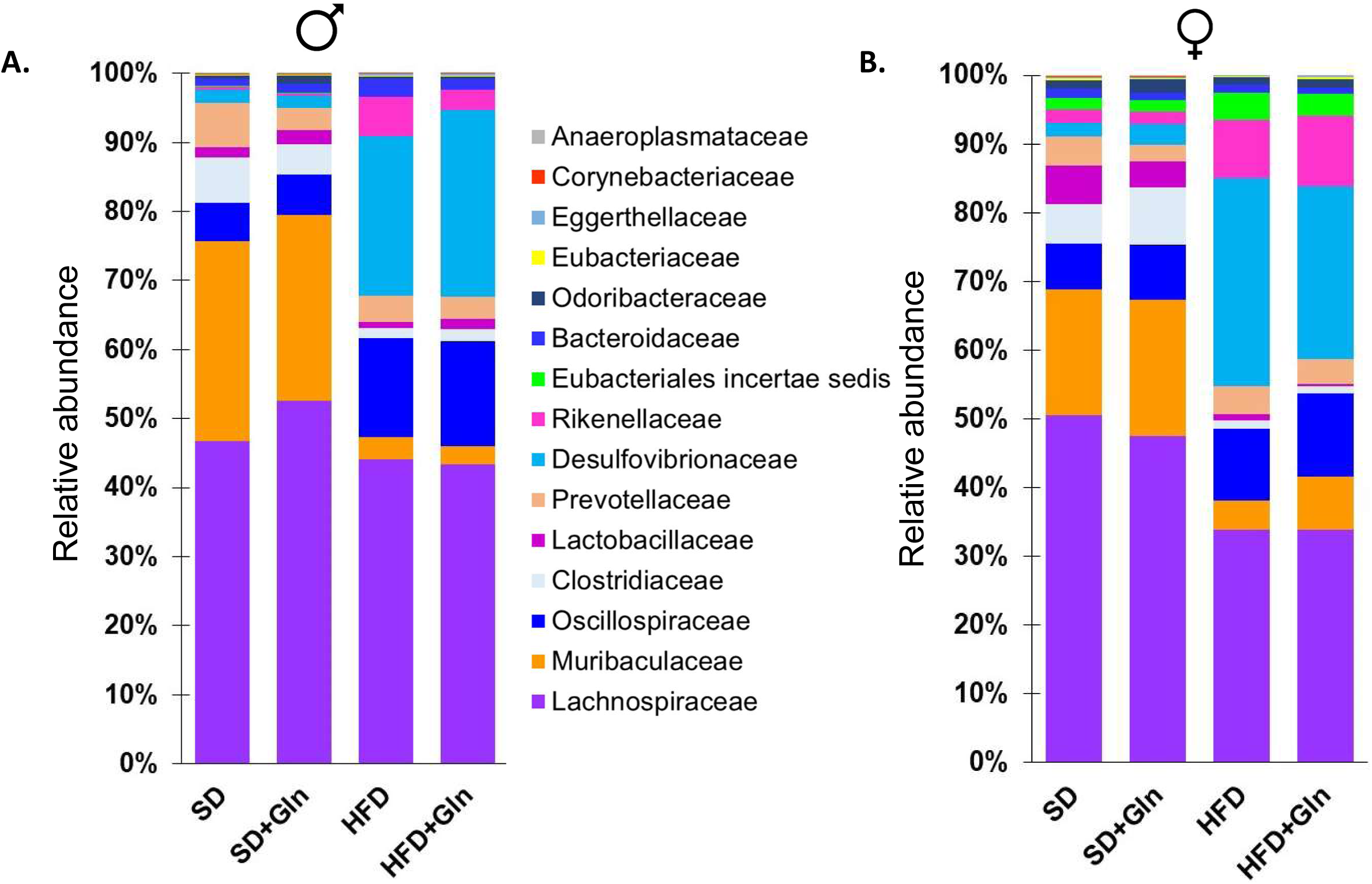
Effects of glutamine (Gln) supplementation on family abundances in the cecal content of male and female mice fed with a standard (SD) or high fat diet (HFD). Family abundance in the cecal content was assessed at week 14 (W14) in male (**A.**) and female (**B.**) mice fed with standard diet (SD) or high fat diet (HFD) and receiving or not glutamine (Gln) in drinking water from W12 to W14. Data are expressed as mean of the percent of relative abundance (n=10-12/group).

**Supplemental Fig. S3:**
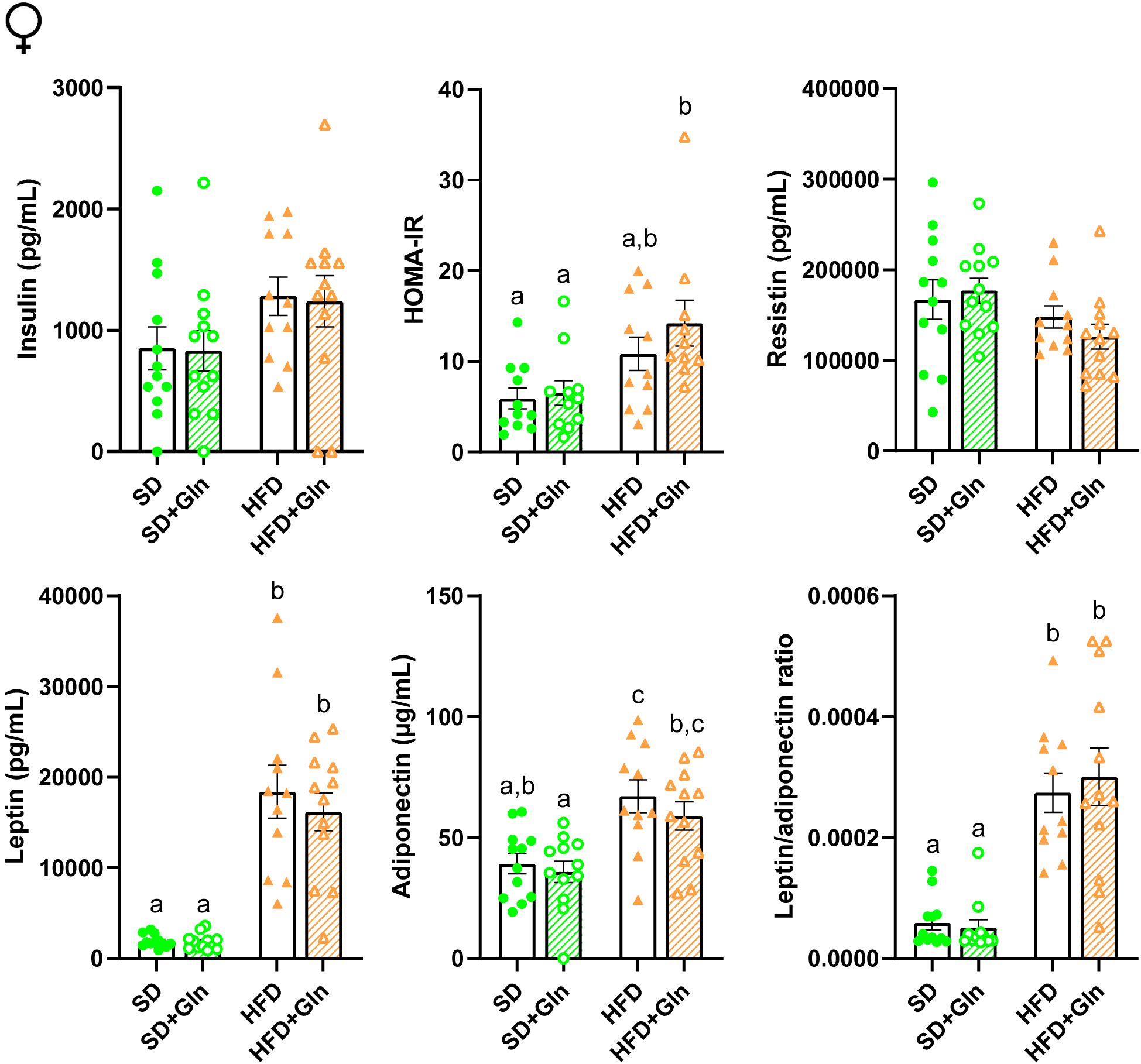
Effects of glutamine (Gln) supplementation on insulinemia, HOMA-IR and plasma adipokine levels in female mice fed with a standard (SD) or high fat diet (HFD). Plasma insulin, resistin, leptin and adiponectin concentrations were assessed at week 14 (W14) in female mice fed with standard diet (SD) or high fat diet (HFD) and receiving or not glutamine (Gln) in drinking water from W12 to W14. The HOMA-IR index was determined by the following equation: fasting serum insulin (μU/mL)LJ×LJfasting plasma glucose (mmol/L)/22.5. Data are expressed as mean ± sem (n=12/group). Values without a common letter differ significantly (p < 0.05).

**Supplemental Fig. S4:**
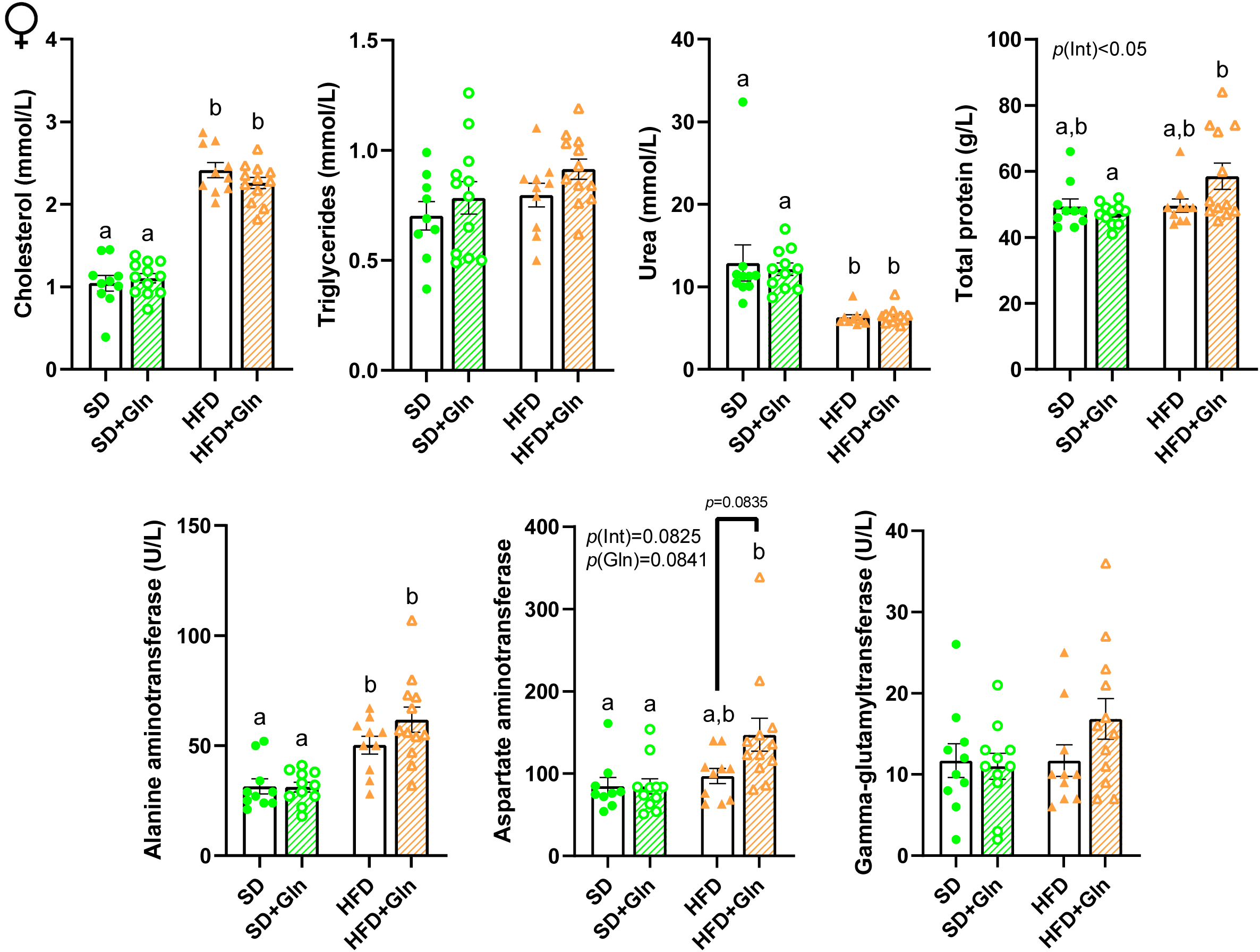
Effects of glutamine (Gln) supplementation on metabolic and liver plasma markers in female mice fed with a standard (SD) or high fat diet (HFD). Cholesterol, triglycerides, urea, total proteins, alanine aminotransferase, aspartate aminotransferase and gamma-glutamyltransferase plasma concentrations were assessed at week 14 (W14) in female mice fed with standard diet (SD) or high fat diet (HFD) and receiving or not glutamine (Gln) in drinking water from W12 to W14. Data are expressed as mean ± sem (n=12/group). Values without a common letter differ significantly (p < 0.05).

**Supplemental Fig. S5:**
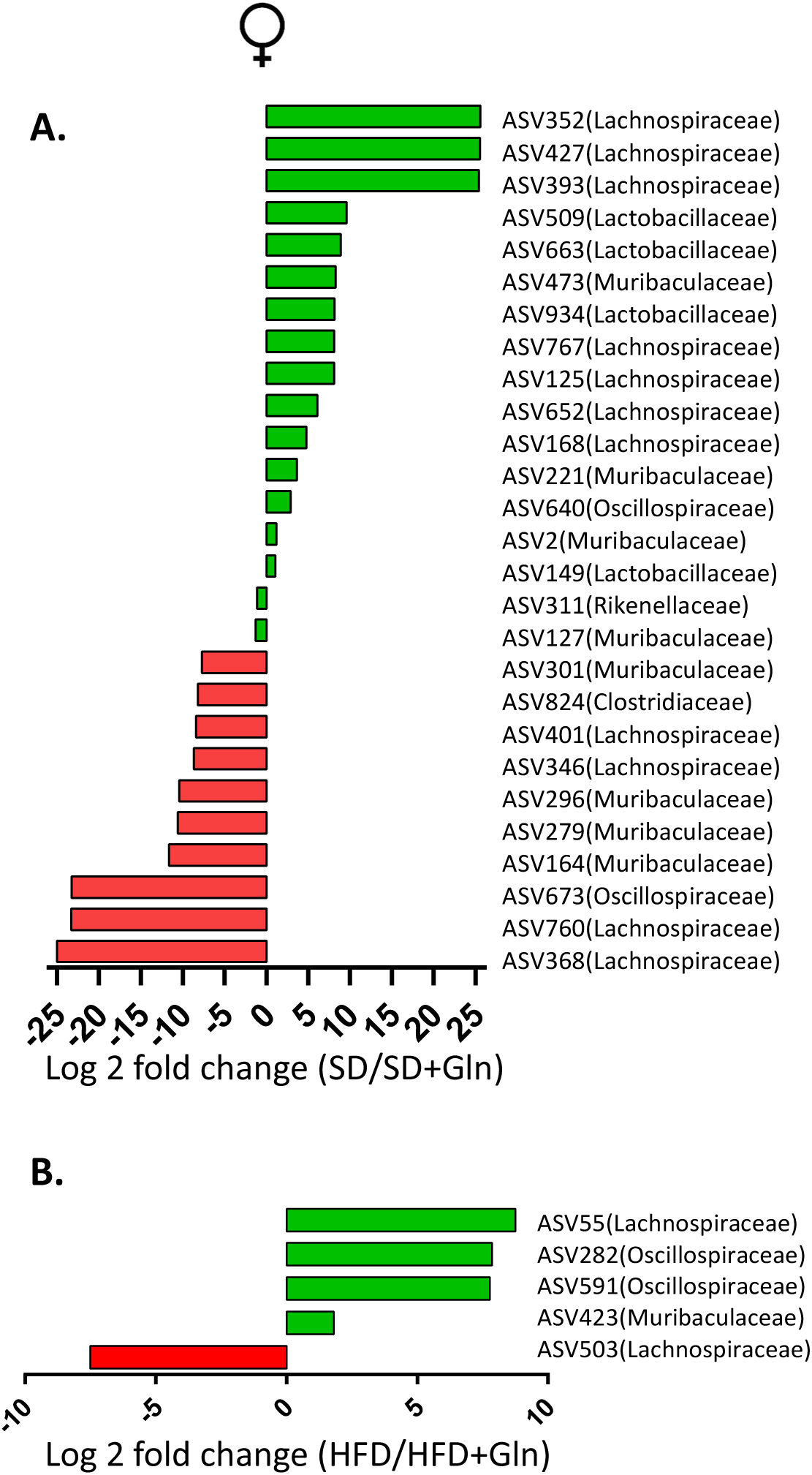
Differentially abundant ASVs in cecal content of female mice fed with standard diet (SD) or high fat diet (HFD) and receiving or not glutamine (Gln). The x-axis indicates the log2(foldchange) (the magnitude of the difference in abundance between the two groups) and the family of ASVs was classified using 16S_REFseq as reference database. (**A.**) Positive log2(FoldChange) value indicates enrichment in the “SD+Gln” treatment group and a negative log2(FoldChange) value indicates enrichment in the “SD” group. (**B.**) Positive log2(FoldChange) value indicates enrichment in the “HFD+Gln” treatment group and a negative log2(FoldChange) value indicates enrichment in the “HFD” group. The ASVs were selected based on effect size (1/2< fold change < 2), adjusted p-value (<0.05) and prevalence (more than 350 copies per sample in at least 25% of the samples).

## References

1. Marques A, Peralta M, Naia A, Loureiro N, de Matos MG. Prevalence of adult overweight and obesity in 20 European countries, 2014. Eur J Public Health. 01 2018;28(2):295LJ300.

2. Ward ZJ, Bleich SN, Cradock AL, Barrett JL, Giles CM, Flax C, et al. Projected U.S. State-Level Prevalence of Adult Obesity and Severe Obesity. N Engl J Med. 19 déc 2019;381(25):2440LJ50.

3. Cani PD, Bibiloni R, Knauf C, Waget A, Neyrinck AM, Delzenne NM, et al. Changes in gut microbiota control metabolic endotoxemia-induced inflammation in high-fat diet-induced obesity and diabetes in mice. Diabetes. juin 2008;57(6):1470LJ81.

4. Cani PD, Amar J, Iglesias MA, Poggi M, Knauf C, Bastelica D, et al. Metabolic endotoxemia initiates obesity and insulin resistance. Diabetes. juill 2007;56(7):1761LJ72.

5. Nascimento JC, Matheus VA, Oliveira RB, Tada SFS, Collares-Buzato CB. High-Fat Diet Induces Disruption of the Tight Junction-Mediated Paracellular Barrier in the Proximal Small Intestine Before the Onset of Type 2 Diabetes and Endotoxemia. Dig Dis Sci. oct 2021;66(10):3359LJ74.

6. de La Serre CB, Ellis CL, Lee J, Hartman AL, Rutledge JC, Raybould HE. Propensity to high-fat diet-induced obesity in rats is associated with changes in the gut microbiota and gut inflammation. Am J Physiol Gastrointest Liver Physiol. août 2010;299(2):G440–448.

7. Bona MD, Torres CH de M, Lima SCVC, Morais AH de A, Lima AÂM, Maciel BLL. Intestinal Barrier Permeability in Obese Individuals with or without Metabolic Syndrome: A Systematic Review. Nutrients. 3 sept 2022;14(17):3649.

8. Teixeira TFS, Souza NCS, Chiarello PG, Franceschini SCC, Bressan J, Ferreira CLLF, et al. Intestinal permeability parameters in obese patients are correlated with metabolic syndrome risk factors. Clin Nutr. oct 2012;31(5):735LJ40.

9. Brignardello J, Morales P, Diaz E, Romero J, Brunser O, Gotteland M. Pilot study: alterations of intestinal microbiota in obese humans are not associated with colonic inflammation or disturbances of barrier function. Aliment Pharmacol Ther. déc 2010;32(11LJ12):1307LJ14.

10. Ott B, Skurk T, Hastreiter L, Lagkouvardos I, Fischer S, Büttner J, et al. Effect of caloric restriction on gut permeability, inflammation markers, and fecal microbiota in obese women. Sci Rep. 20 sept 2017;7(1):11955.

11. Damms-Machado A, Louis S, Schnitzer A, Volynets V, Rings A, Basrai M, et al. Gut permeability is related to body weight, fatty liver disease, and insulin resistance in obese individuals undergoing weight reduction. Am J Clin Nutr. janv 2017;105(1):127LJ35.

12. Koutoukidis DA, Jebb SA, Zimmerman M, Otunla A, Henry JA, Ferrey A, et al. The association of weight loss with changes in the gut microbiota diversity, composition, and intestinal permeability: a systematic review and meta-analysis. Gut Microbes. 2022;14(1):2020068.

13. Petrus P, Lecoutre S, Dollet L, Wiel C, Sulen A, Gao H, et al. Glutamine Links Obesity to Inflammation in Human White Adipose Tissue. Cell Metab. 4 févr 2020;31(2):375–390.e11.

14. Achamrah N, Déchelotte P, Coëffier M. Glutamine and the regulation of intestinal permeability: from bench to bedside. Curr Opin Clin Nutr Metab Care. 2017;20(1):86LJ91.

15. Beutheu S, Ghouzali I, Galas L, Déchelotte P, Coëffier M. Glutamine and arginine improve permeability and tight junction protein expression in methotrexate-treated Caco-2 cells. Clin Nutr. oct 2013;32(5):863LJ9.

16. Beutheu S, Ouelaa W, Guérin C, Belmonte L, Aziz M, Tennoune N, et al. Glutamine supplementation, but not combined glutamine and arginine supplementation, improves gut barrier function during chemotherapy-induced intestinal mucositis in rats. Clin Nutr. août 2014;33(4):694LJ701.

17. Ghouzali I, Lemaitre C, Bahlouli W, Azhar S, Bôle-Feysot C, Meleine M, et al. Targeting immunoproteasome and glutamine supplementation prevent intestinal hyperpermeability. Biochim Biophys Acta. 2017;1861(1 Pt A):3278LJ88.

18. Bertrand J, Ghouzali I, Guérin C, Bôle-Feysot C, Gouteux M, Déchelotte P, et al. Glutamine Restores Tight Junction Protein Claudin-1 Expression in Colonic Mucosa of Patients With Diarrhea-Predominant Irritable Bowel Syndrome. JPEN J Parenter Enteral Nutr. 2016;40(8):1170LJ6.

19. Zhou Q, Verne ML, Fields JZ, Lefante JJ, Basra S, Salameh H, et al. Randomised placebo-controlled trial of dietary glutamine supplements for postinfectious irritable bowel syndrome. Gut. 2019;68(6):996LJ1002.

20. Choi K, Lee SS, Oh SJ, Lim SY, Lim SY, Jeon WK, et al. The effect of oral glutamine on 5-fluorouracil/leucovorin-induced mucositis/stomatitis assessed by intestinal permeability test. Clin Nutr. févr 2007;26(1):57LJ62.

21. Opara EC, Petro A, Tevrizian A, Feinglos MN, Surwit RS. L-glutamine supplementation of a high fat diet reduces body weight and attenuates hyperglycemia and hyperinsulinemia in C57BL/6J mice. J Nutr. janv 1996;126(1):273LJ9.

22. Laviano A, Molfino A, Lacaria MT, Canelli A, De Leo S, Preziosa I, et al. Glutamine supplementation favors weight loss in nondieting obese female patients. A pilot study. Eur J Clin Nutr. nov 2014;68(11):1264LJ6.

23. Abboud KY, Reis SK, Martelli ME, Zordão OP, Tannihão F, de Souza AZZ, et al. Oral Glutamine Supplementation Reduces Obesity, Pro-Inflammatory Markers, and Improves Insulin Sensitivity in DIO Wistar Rats and Reduces Waist Circumference in Overweight and Obese Humans. Nutrients. 1 mars 2019;11(3).

24. de Souza AZZ, Zambom AZ, Abboud KY, Reis SK, Tannihão F, Guadagnini D, et al. Oral supplementation with L-glutamine alters gut microbiota of obese and overweight adults: A pilot study. Nutrition. juin 2015;31(6):884LJ9.

25. Lefebvre C, Tiffay A, Breemeersch CE, Dreux V, Bôle-Feysot C, Guérin C, et al. Sex-dependent effects of a high fat diet on metabolic disorders, intestinal barrier function and gut microbiota in mouse. Sci Rep. 27 août 2024;14(1):19835.

26. Medrikova D, Jilkova ZM, Bardova K, Janovska P, Rossmeisl M, Kopecky J. Sex differences during the course of diet-induced obesity in mice: adipose tissue expandability and glycemic control. Int J Obes (Lond). févr 2012;36(2):262LJ72.

27. Elzinga SE, Savelieff MG, O’Brien PD, Mendelson FE, Hayes JM, Feldman EL. Sex differences in insulin resistance, but not peripheral neuropathy, in a diet-induced prediabetes mouse model. Dis Model Mech. 1 avr 2021;14(4):dmm048909.

28. Peng C, Xu X, Li Y, Li X, Yang X, Chen H, et al. Sex-specific association between the gut microbiome and high-fat diet-induced metabolic disorders in mice. Biol Sex Differ. 20 janv 2020;11(1):5.

29. McMurdie PJ, Holmes S. phyloseq: an R package for reproducible interactive analysis and graphics of microbiome census data. PLoS One. 2013;8(4):e61217.

30. Wang B, Wang L, Wang H, Dai H, Lu X, Lee YK, et al. Targeting the Gut Microbiota for Remediating Obesity and Related Metabolic Disorders. J Nutr. 1 juill 2021;151(7):1703LJ16.

31. L’Huillier C, Jarbeau M, Achamrah N, Belmonte L, Amamou A, Nobis S, et al. Glutamine, but not Branched-Chain Amino Acids, Restores Intestinal Barrier Function during Activity-Based Anorexia. Nutrients. 15 juin 2019;11(6).

32. Bakalar B, Duska F, Pachl J, Fric M, Otahal M, Pazout J, et al. Parenterally administered dipeptide alanyl-glutamine prevents worsening of insulin sensitivity in multiple-trauma patients. Crit Care Med. févr 2006;34(2):381LJ6.

33. Déchelotte P, Hasselmann M, Cynober L, Allaouchiche B, Coëffier M, Hecketsweiler B, et al. L-alanyl-L-glutamine dipeptide-supplemented total parenteral nutrition reduces infectious complications and glucose intolerance in critically ill patients: the French controlled, randomized, double-blind, multicenter study. Crit Care Med. mars 2006;34(3):598LJ604.

34. Palomo-Buitrago ME, Sabater-Masdeu M, Moreno-Navarrete JM, Caballano-Infantes E, Arnoriaga-Rodríguez M, Coll C, et al. Glutamate interactions with obesity, insulin resistance, cognition and gut microbiota composition. Acta Diabetol. mai 2019;56(5):569LJ79.

35. Lin XB, Dieleman LA, Ketabi A, Bibova I, Sawyer MB, Xue H, et al. Irinotecan (CPT-11) chemotherapy alters intestinal microbiota in tumour bearing rats. PLoS One. 2012;7(7):e39764.

36. Jegatheesan P, Beutheu S, Ventura G, Sarfati G, Nubret E, Kapel N, et al. Effect of specific amino acids on hepatic lipid metabolism in fructose-induced non-alcoholic fatty liver disease. Clin Nutr. févr 2016;35(1):175LJ82.

37. Dai ZL, Li XL, Xi PB, Zhang J, Wu G, Zhu WY. L-Glutamine regulates amino acid utilization by intestinal bacteria. Amino Acids. sept 2013;45(3):501LJ12.

38. Bu Y, Liu Y, Liu Y, Cao J, Zhang Z, Yi H. Protective Effects of Bacteriocin-Producing Lactiplantibacillus plantarum on Intestinal Barrier of Mice. Nutrients. 10 août 2023;15(16):3518.

39. Xie A, Song J, Lu S, Liu Y, Tang L, Wen S. Influence of Diet on the Effect of the Probiotic Lactobacillus paracasei in Rats Suffering From Allergic Asthma. Front Microbiol. 2021;12:737622.

40. Pereira FC, Wasmund K, Cobankovic I, Jehmlich N, Herbold CW, Lee KS, et al. Rational design of a microbial consortium of mucosal sugar utilizers reduces Clostridiodes difficile colonization. Nat Commun. 9 oct 2020;11(1):5104.

41. Wu Z, Du Z, Tian Y, Liu M, Zhu K, Zhao Y, et al. Inulin accelerates weight loss in obese mice by regulating gut microbiota and serum metabolites. Front Nutr. 2022;9:980382.

42. Wu Z, Zhang M, Deng Y, Zhou G, Yang M, Wang H. Alterations of gut microbiome and metabolism induced by inulin associated with weight loss in obese female mice. Int J Food Sci Nutr. sept 2023;74(5):606LJ20.

43. Salazar-Jaramillo L, de la Cuesta-Zuluaga J, Chica LA, Cadavid M, Ley RE, Reyes A, et al. Gut microbiome diversity within Clostridia is negatively associated with human obesity. mSystems. 20 août 2024;9(8):e0062724.

44. Hao J, Zhang J, Wu T. Fucoxanthin extract ameliorates obesity associated with modulation of bile acid metabolism and gut microbiota in high-fat-diet fed mice. Eur J Nutr. févr 2024;63(1):231LJ42.

